# PyPE_RESP: A tool to facilitate and standardize derivation of RESP charges

**DOI:** 10.1101/2025.01.07.631713

**Authors:** Marco Lapsien, Michele Bonus, Lianne Gahan, Adélaïde Raguin, Holger Gohlke

## Abstract

We introduce PyPE_RESP, a tool to facilitate and standardize partial atomic charge derivation using the RESP approach. PyPE_RESP builds upon the open-source Python package RDKit for chemoinformatics and the AMBER suite for molecular simulations. PyPE_RESP provides an easy setup of multi-conformer and multi-molecule Restrained Electrostatic Potential (RESP) fitting while allowing a comprehensive definition of charge constraint groups through multiple methods. As a command line tool, PyPE_RESP can be integrated into batch processes. The software enables the derivation of partial atomic charges for additive and polarizable force fields. It outputs constraint group and non-constraint group charges to give an immediate overview of the fit result. PyPE_RESP will be distributed with AmberTools and compatible with most computing platforms.

**Table of Contents Graphic:** 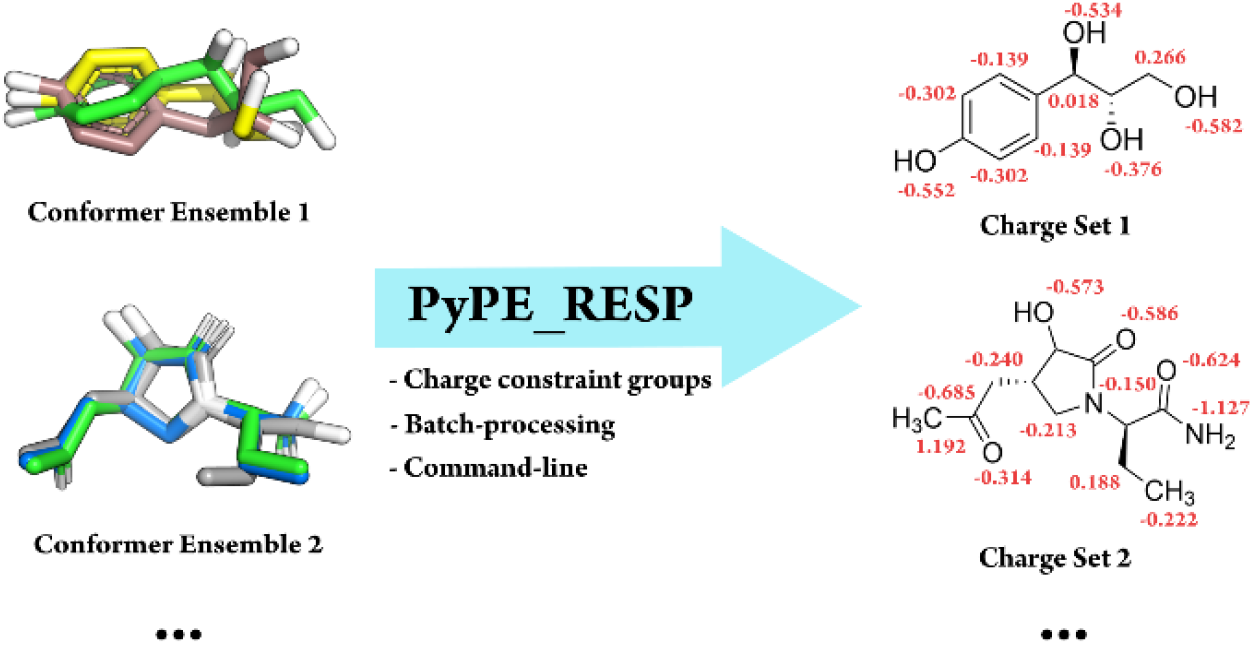

## Introduction

The accurate representation of electrostatic interactions is essential for developing, extending, and applying empirical force fields used in molecular dynamics simulations.^1^ Considering the prominent use of wide-spread additive force fields such as from the AMBER family^2,3,4^ or CHARMM family,^5,6^ and the on-going development of polarizable force fields such as the atomic multipole-optimized energetics for biomolecular applications (AMOEBA) force field,^7,8^ this statement holds regardless of the type of force field being developed. Although various methods for deriving partial atomic charges have been described,^9^ the restrained electrostatic potential (RESP) approach is still the main charge derivation method used in modern AMBER force fields.^10,11^ This can be attributed to its capability to reproduce the quantum mechanical molecular electrostatic potential (ESP) while producing chemically sensible charge distributions.^12^ Though RESP was initially developed to obtain fixed atom-centered partial charges for additive force fields, it has been further evolved into the RESP-ind and RESP-perm models implemented in PyRESP^13^ with induced and permanent dipole moments for polarizable force fields. A major drawback of the RESP method lies in its setup requirements, as multiple preparation steps must be carried out before obtaining atomic charges for the system of interest. These steps include:

i. Extraction of electrostatic potential (ESP) information from Gaussian^14^ output files (*via* espgen^15^)
ii. Generation of input files for PyRESP (*via* Py-RESP_GEN^16^)
iii. Insertion of charge constraint and atom equivalency information into PyRESP input files.

While tasks i) and ii) can be automated by dedicated software, task iii) either requires manual editing of input files or creating custom scripts to adhere to the RESP input file format. It additionally requires the manual definition of charge constraint groups (CCGs), meaning a group of atoms whose charge is fixed during fitting, by referencing the respective atom indices. The latter can be tedious and error-prone when PyRESP needs to be invoked for multiple molecules. Further complications arise when multiple conformers of a given molecule are considered. Currently, Py-RESP_GEN is only capable of producing PyRESP input files for one structure at a time, which therefore involves additional file management by the user to obtain a concatenated PyRESP input file. This also applies to the ESP information, which must be concatenated into a single file. Finally, the obtained RESP charges must be manually input into a structure file (e.g., mol2 format), as charges will only be written to a parameter file created by PyRESP. These concerns have already been tackled to some extent by the development of the RESP ESP charge derive (R.E.D.) server.^17^ Although the R.E.D. server covers tasks i) and ii) well and offers various functionalities, such as integrated geometry optimization and ESP calculation using Gaussian, its syntax for defining CCGs can be overwhelming if many structures need to be parameterized. This is also reflected in the server’s naming convention for output files, which can impede the identification of a desired molecular fragment. Additionally, the use of an external server always carries a risk of experiencing periods of downtime and requires uploading research data. Furthermore, antechamber^15^, a tool within the AMBER molecular dynamics suite, is capable of rapidly deriving RESP charges, but is limited to single conformer fitting. Therefore, the conformational bias of RESP charges is neglected in this approach^18^.

To overcome these issues, we developed PyPE_RESP, a Python-based workflow to conveniently and systematically execute multi-conformer and multi-molecule RESP fitting within the realm of AMBER, using Gaussian for ESP computation. PyPE_RESP will be distributed under a GPL license together with Amber-Tools^15^ (https://ambermd.org/AmberTools.php).

### Workflow description

The PyPE_RESP workflow minimizes the user’s manual work while providing flexible setups and reliable and reproducible results of charge fitting. A flowchart of the main steps of PyPE_RESP is presented in **Figure 1**. PyPE_RESP builds upon existing tools (antechamber^15^, espgen, PyRESP_GEN^16^, and PyRESP^13^) and manages their in- and outputs, enabling it to be adaptable towards specific properties of each system to be parametrized and forming an “all-in-one” solution for RESP charge derivation. The sole input files required are the Gaussian output files containing the ESP information for each conformer of each molecule. These files must be gathered in a dedicated directory. All other specifications for the RESP run are managed by flags submitted *via* the PyPE_RESP command line interface (CLI) or a PyPE_RESP input file. The latter is convenient for the repetitive processing of multiple molecules. Most flags in the CLI utilize PyPE_RESP’s inherent enumeration of CCGs, which is illustrated in **Figure 2**.

**Figure 1:**
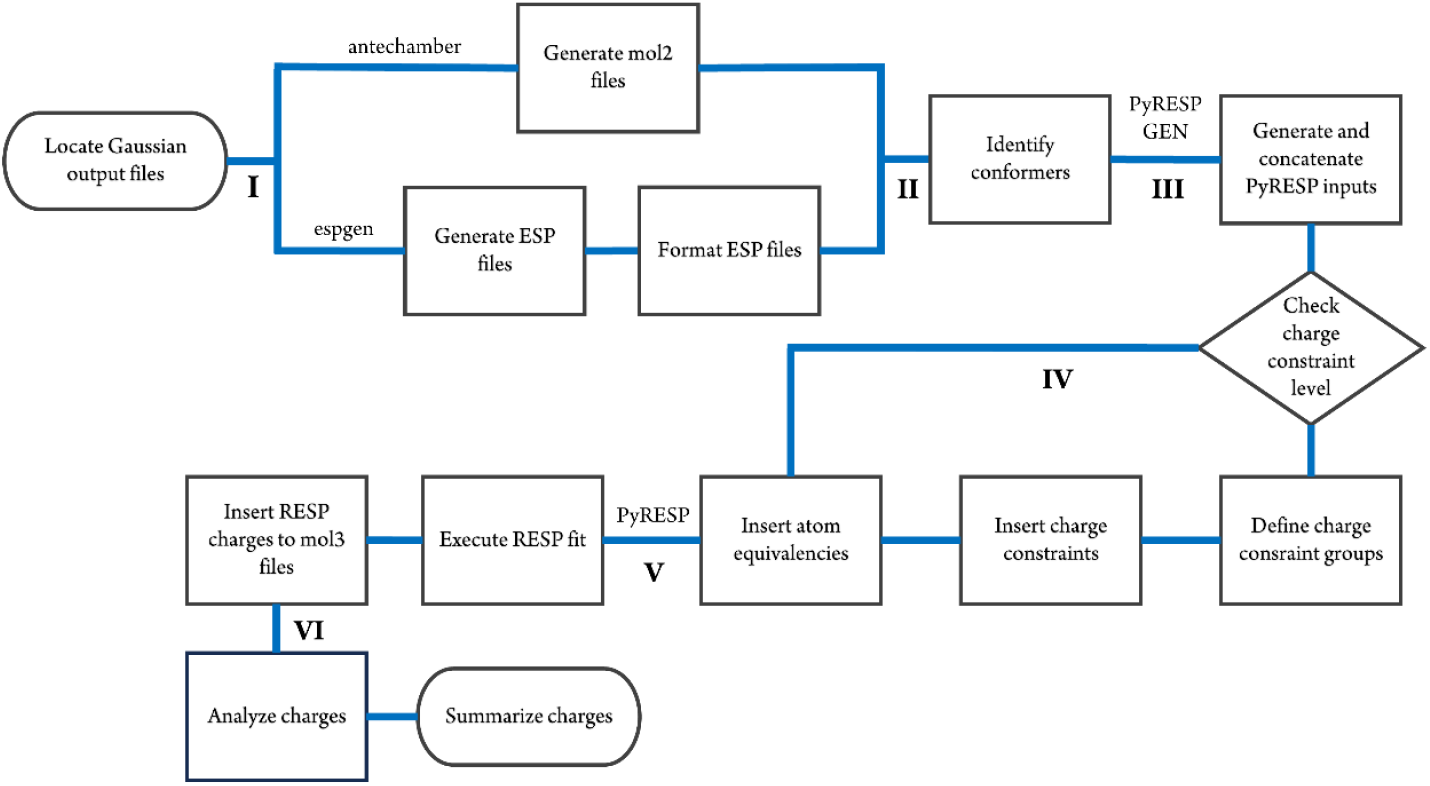
Flowchart highlighting the most important steps in the PyPE_RESP workflow. After locating each Gaussian output file, mol2 and ESP files are created simultaneously (I). Then, the number of conformers per unique molecule is identified by comparing the International Chemical Identifier (InChI) of each molecule (II). First-stage and second-stage PyRESP input files are created by executing PyRESP_GEN and subsequently concatenated to yield two PyRESP input files (III). Depending on the charge constraint level chosen, CCGs are defined based on the information supplied on the command line and appended to the PyRESP input files. The atom equivalency information (i.e., declaring that atoms of all conformers are treated equally) is also appended to the PyRESP input files (IV). Next, PyRESP is executed to start the fitting process (V). The resulting partial atomic charges are written to a newly generated mol3 file for each molecule together with the molecule’s connectivity points, meaning atoms connected to, but not part of a constraint group, determined by PyPE_RESP. The derived charges are analyzed based on the mol3 files by utilizing the atom indices of each constraint group to match the corresponding atom charges (VI). Additionally, the sum of charges for each constraint group and the sum of charges of each atom not belonging to the constraint group are calculated. This information is written in a log file and a csv file.

**Figure 2:**
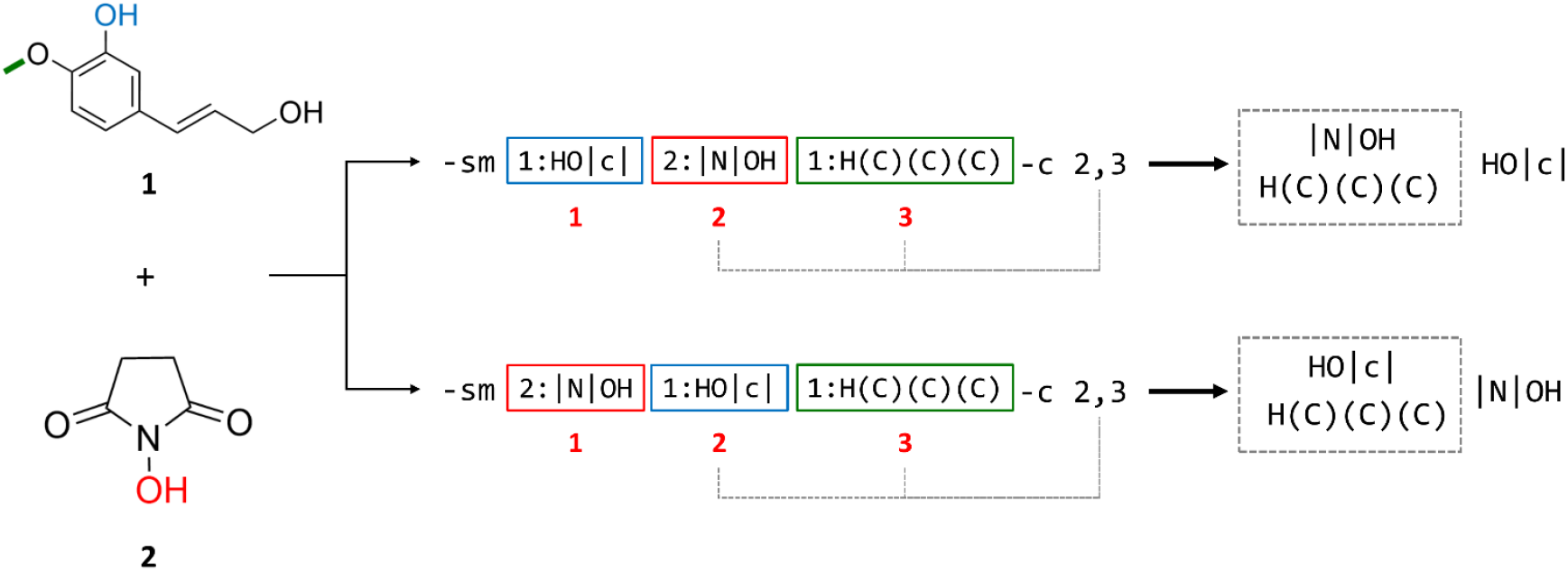
Illustration of PyPE_RESP’s inherent constraint group enumeration on arbitrary constraints. An intermolecular CCG between the methyl group of compound **1** (green) and the hydroxyl group of compound **2** (red) is defined. Additionally, an intramolecular CCG comprising the hydroxyl group of compound **1** (blue) is defined. Each constraint group is identified using SMARTS substructure matching utilizing the `-sm (--smarts_match)` flag. Constraint groups are indexed (starting from 1) based on the order they are defined under the `-sm (or -cl` for atom indexing) flag. The assigned indices are then used by the `-c (--combine)` flag to determine, which of the constraint groups are used to define intermolecular charge constraints (dashed box). Changing the order in which CCGs are defined (e.g., swapping the positions of CCGs at positions 1 and 2) without changing index references in the -c flag leads to altered targeting behavior. Here, this leads to the faulty definition of an intermolecular CCG between the hydroxyl and methyl groups of compound **1**, while the hydroxyl group of compound **2** is used to define an intramolecular CCG.

### Definition of charge constraint groups

During the electrostatic parametrization of fragments derived from large molecules, they are often equipped with a capping group that mimics the chemical environment in the assembled molecule. Since these capping groups are only attached temporarily, charge constraints are applied to account for their removal after partial atomic charges have been derived. PyPE_RESP indexes each CCG (starting from 1) in the order they are defined under the respective CLI flag. This index can be used to specify if a CCG is part of an intermolecular group constraint (e.g., for considering capping groups). Whether group constraints are considered during the RESP fit is managed by setting the constraint level using the `--constraints` flag. It can be set to ignore charge constraints (0) or allow intramolecular charge constraints (1) and/or intermolecular charge constraints (2). Consequently, the definition of CCGs is only allowed if at least intramolecular constraints are chosen.

CCGs can be defined in two ways. First, by utilizing the `--con_list` flag to supply groups of atom indices to be constrained. The atom indices can be taken directly from the Gaussian output files that PyPE_RESP uses as inputs. Still, a visual inspection of the molecule’s three-dimensional structure is most likely required to determine all indices of atoms of a group constraint. While visualization software exists that can read Gaussian output files, common molecule visualization software, such as VMD^19^ or PyMOL,^20^ cannot do this without further modification. To overcome this, PyPE_RESP allows to carry out a conversion run if the `--mol2_only` flag is supplied. In this case, PyPE_RESP will generate mol2 files for each Gaussian output file found using antechamber and exit afterward. Second, defining CCGs is based on matching substructures of molecules generated from SMARTS patterns by utilizing the RDKit Open-source cheminformatics package.^21^ These can be supplied under the `--smarts_match` flag (**Figure 2**). Both `--con_list` and `--smarts_match` follow the same syntax where the molecule to assign the CCG to is specified, followed by a colon and the SMARTS pattern to be used for substructure matching, or the comma-separated atom indices. Molecules are indexed starting from 1 and can be referenced by their index to define CCGs. Indexing is either determined automatically based on the ascending alphabetical order of filenames or by specifying wildcard patterns matching the respective filenames, which will be resolved using Python’s glob module. The `--smarts_match` and `--con_list flags` are mutually exclusive, i.e., all CCGs must be either defined by SMARTS patterns or atom indices. Additionally, PyPE_RESP only accepts unambiguous SMARTS patterns and raises an error if multiple substructure matches are found for a given pattern.

The use of SMARTS patterns often requires less initial setup than defining CCGs by atom indices but can create long and complex character sequences to match a specific functional group. To verify whether a pattern matches one or multiple substructures the `--test_smarts` flag can be utilized. It returns the number of substructure matches per molecule, along with the atom indices corresponding to each match for every SMARTS pattern passed as an argument. If a pattern is found to be ambiguous, a warning is displayed. An incorrectly specified SMARTS string will force the program to exit.

SMARTS patterns often include atoms that are required to ensure unambiguity but are not meant to be part of the final CCG. To neglect these atoms later during fitting, “SMARTS slicing” can be applied by enclosing the parts of the SMARTS pattern to be neglected in pipe symbols (**Figure 2**). PyPE_RESP recognizes these symbols and removes the atom indices corresponding to the enclosed section after the full pattern has been used for substructure matching.

### Setting fit properties

Besides defining single CCGs, one needs to consider how intermolecular charge constraints are defined. This is done with the –c (`--combine)` flag, which specifies which CCGs are used to define intermolecular charge constraints. The `--combine` flag identifies CCGs from `--smarts_match` or `--con_list` definitions based on their order-dependent indices (**Figure 2**). Each CCG index not referenced under the `--combine` flag will be used to define an intramolecular CCG.

The assignments of constraint group and molecular charges are another crucial step in RESP charge derivation. While molecular charges do not need to be defined as they are taken from the respective Gaussian input file, constraint group charges are set to zero by default, but can be assigned by specifying the group charge during the definition of the respective constraint group (see Examples section).

Besides the traditional RESP method, PyPE_RESP also allows using the RESP-ind and RESP-perm models that are integrated into the PyRESP software. The associated flags `(--ptype` and `--polariz`) are identical to those used in PyRESP and PyRESP_GEN. These flags do not need to be specified if the usual point-charge model (“chg”) is chosen.

PyRESP execution can be prohibited by supplying the `--no_pyresp` flag, which stops PyPE_RESP after the generation of PyRESP input files, to allow for analysis by the user. PyPE_RESP also provides insights and statistics for each run carried out. A log file is created that documents the key steps during the generation of the PyRESP input files. It includes the number of unique molecules found and the number of conformers per molecule detected. Moreover, a second log file is created after the successful execution of PyRESP, which contains the fit statistics provided by PyRESP and highlights the sum of charges for each CCG, the sum of charges of all atoms that do not belong to the respective constraint group, and the difference between these two sums. This information is also stored in a csv file to facilitate integration into data analysis software for further inspection.

### Examples

Each one of the following example cases (**Figure 3**) will be carried out as a multi-conformer RESP fit. Refer to the Supporting Information (SI) for details on conformer generation. The described examples focus on using the PyPE_RESP CLI instead of PyPE_RESP input files. Alternatively, input files, including Gaussian output files and PyPE_RESP input files, for each example are provided in the SI.

**Figure 3:**
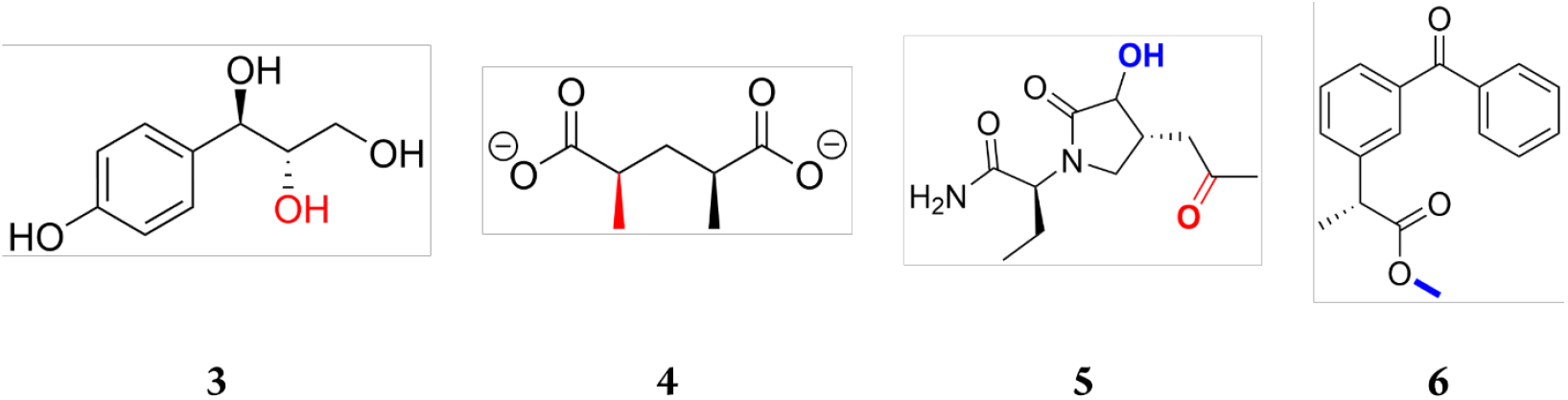
Example compounds to illustrate the use of PyPE_RESP. **3**: (1*R*,2*S*)-1-(4-hydroxyphenyl)propane-1,2,3-triol; **4**: (2*R*,4*S*)-2,4-dimethylpentanedioate; **5**: (2*S*)-2-((4*R*)-3-hydroxy-2-oxo-4-(2-oxopropyl)pyrrolidin-1-yl)butanamide; **6**: (2*R*)-methyl-2-(3-benzoylphenyl)propanoate. Intramolecular CCGs are highlighted in red, whereas groups that are part of an intermolecular charge constraint are shown in blue.

CASE A: Using SMARTS patterns to define CCGs.

The benefits of SMARTS slicing to define an intramolecular CCG can be illustrated using the hydroxyl group bound to the carbon atom at position two of molecule **3** (**Figure 3**). Since this compound possesses multiple hydroxyl groups, the unambiguous SMARTS pattern to identify the group of interest must include the neighboring aliphatic carbon atom as well as the aromatic carbon attached to it. This results in the following SMARTS pattern: HOCCc. The three carbon atoms in this sequence are not meant to be part of the final CCG and, therefore, must be neglected during fitting using SMARTS slicing. To achieve this, the three carbon atoms must be enclosed in pipe symbols. The whole pattern must also be enclosed in quotation marks to prevent interpretation by the shell. Thus, the full command to identify the 2-OH group for an intramolecular CCG is: `pype-resp.py –constraints 1 --smarts_match 1:”HO|CCc|”`.

Notably, the hydrogen atom is not enclosed in square brackets, unlike required by the SMARTS language convention. PyPE_RESP adds the brackets automatically, although they will be recognized if added by the user. Identification of the target molecule by its index (1) was chosen here for convenience as it is the only molecule to be parametrized. A wildcard pattern to identify the molecule would also work.

CASE B: Defining intramolecular CCGs using atom indices and the RESP-ind model

CCG definition *via* SMARTS patterns can become difficult when dealing with symmetric molecules, and defining them by the respective atom indices can then be easier. For example, this occurs when targeting one of the two methyl groups of molecule **4** (**Figure 3**) to define an intramolecular CCG. A SMARTS pattern will always also match the molecule’s other methyl group and therefore is ambiguous. PyPE_RESP offers two possibilities to solve this issue. One option is to carry out a test run by supplying the `--test_smarts` flag with the SMARTS pattern C(H)(H)(H), which retrieves the atom indices corresponding to the two methyl groups. The indices of one of these groups can then be used to define a CCG using the `--con_list` flag in a production run. Alternatively, one can carry out a PyPE_RESP run with the `--mol2_only` flag to generate mol2 files, which can then be visualized to obtain the atom indices of the methyl group atoms. Both approaches reveal the atoms with indices 1, 12, 13, and 14 to be part of the target functional group. Atomic partial charges for the compound can be obtained using the RESP-ind model, for which the “ind” keyword must be supplied under the `--ptype` flag. The use of the RESP-ind and RESP-perm models requires information about individual atomic polarizabilities. The absolute or relative path to a file containing this information can be supplied under the `--polariz` flag (an example is provided in the Supporting Information). Finally, the charge of the constraint group for the methyl moiety must be set to −1 by adding the colon-separated charge after the definition of the constraint group. The full command is: `pype-resp.py --constraints 1 --con_list 1:1,12,13,14:-1 --ptype ind --polariz Polarizability_info.txt`

CASE C: Defining intermolecular constraints by SMARTS matching and wildcard patterns

If multiple molecules need to be parametrized, it is reasonable to identify them *via* wildcard patterns instead of their CCG indices to avoid mismatches. The `--combine` flag is required to manage which of the defined constraint groups will form intermolecular CCGs. Any constraint group not mentioned under the flag is used to define an intramolecular CCG instead. Taking molecule **5** and molecule **6** (**Figure 3**), the hydroxyl group attached to the pyrrolidine ring of **5** should be constrained together with the methyl group of **6**. The carbonyl carbon and carbonyl oxygen of **5** should additionally form an intramolecular CCG, whose charge should be set to −1. This results in the definition of three CCGs, with two being constrained together. The typical command that achieves this is: `pype-resp.py --constraints 2 –-smarts_match butanamide:”OH”`

`propanoate:”C(H)(H)(H)|O|”`

`butanamide:”|[C;H3]`|`C=O”:-1 --combine 1,2:0`

As highlighted in **Figure 2**, the order in which arguments are supplied under the `--smarts_match` flag is important, since in this case `--combine` forces the first and second SMARTS patterns specified to form an intermolecular CCG. This example also illustrates how the charge for intermolecular constraint groups can be set. Similar to intramolecular CCGs, the charge is set after specifying which constraint groups are part of the respective intermolecular CCG. As mentioned above, if the charge is supposed to be zero, no specification is needed, and it is only specified here for illustration purposes. Finally, note that wildcard patterns identifying the respective molecules do not contain asterisks. These are added automatically by PyPE_RESP during the globing process.

## Discussion

We present PyPE_RESP to facilitate and standardize the derivation of atomic partial charges following the RESP procedure. PyPE_RESP uses the RDKit opensource chemoinformatics package and those built into PyPE_RESP of AmberTools to derive charges for additive or polarizable force fields, while providing multiple means to define CCGs in non-trivial cases. PyPE_RESP improves the reproducibility of parameterization tasks, which would otherwise be carried out by hand, and provides informative post-run statistics that allow evaluating the fitting process. While most of these features are also implemented in the R.E.D. server, PyPE_RESP offers the benefit of being executed locally. This circumvents depending on an external server, which eliminates issues arising from downtime periods as well as the need to upload scientific data, and makes the resubmission of jobs easier if minor mistakes in the command or input file were detected in a prior run.

We acknowledge that the syntax required by PyPE_RESP can be overwhelming initially. However, in the context of force field design, or comprehensive molecular design and analysis studies, where numerous compounds must be parameterized, the benefits of PyPE_RESP’s systematic approach stand out. Similarly, creating SMARTS patterns to target components of complex molecules such as multi-ring systems can be troublesome, e.g., if they are intertwined. In such a case, an inexperienced user may need to go through a lengthy trial-and-error process to match the desired substructure. Such scenarios can be prevented, however, by falling back to the definition of CCGs *via* atom indices, further highlighting that PyPE_RESP’s functionality can be adapted according to system complexity. Although PyPE_RESP offers a range of options to cover the needs of most RESP charge derivation procedures, it does not yet account for niche cases, where additional fine-tuning of parameters or reorientation of molecules before fitting is required.

Finally, PyPE_RESP is compatible with most operating systems. This includes common Linux distributions, MacOS, and Windows 11 by using the Windows Subsystem for Linux (WSL).

## ASSOCIATED CONTENT

### Supporting Information

Details on the generation of conformer ensembles and ESP calculation for each compound discussed in the Examples section are described in the SI.

Additionally, the Gaussian output files per conformer for each molecule discussed in the Examples section, along with PyPE_RESP input files and all files generated by PyPE_RESP for each example case, are provided.

This material is available free of charge *via* the Internet at http://pubs.acs.org

## AUTHOR INFORMATION

## Author Contributions

HG designed the study; ML wrote the software code; MB and LG supported software design; ML, AR, and HG wrote the manuscript; MB and LG revised the manuscript; HG and AR secured funding.

## Notes

The authors declare no competing financial interest.

## ACKNOWLEDGMENTS

We are grateful for the computational infrastructure and support provided by the “Zentrum für Informations-und Medientechnologie” (ZIM) at Heinrich Heine University Düsseldorf and the computing time provided by the John von Neumann Institute for Computing (NIC) to HG on the supercomputer JUWELS at Jülich Supercomputing Centre (JSC) (user IDs: VSK33, lignin). We acknowledge the developers of AMBER and the RDKit cheminformatics package for making their software available publicly.

## ABBREVIATIONS

AMBER: Assisted Model Building with Energy Refinement
AMOEBA: Atomic Multipole Optimized Energetics for Biomolecular Applications
CCG: Charge constraint group
CLI: Command Line Interface
ESP: Electrostatic Potential
InChI: International Chemical Identifier
R.E.D.: RESP ESP charge Derive
RESP: Restrained Electrostatic Potential
SMARTS: SMILES Arbitrary Target Specification
WSL: Windows Subsystem for Linux.

