## Supporting Information for "PyPE_RESP: A tool to facilitate and standardize derivation of RESP charges"

### Supplemental methods

#### Conformer generation

Three-dimensional coordinates were generated and stored in a PDB file for a given molecule to generate conformers using the RDKit open-source cheminformatics package[1] (random seed 0xf00d). The resulting PDB files were converted to the xyz file format through the Open Babel[2] command line interface, and a subsequent geometry optimization was performed by using the ANCopt tool built into XTB.[3] The energy and gradient convergence were set to $5\cdot10^{-6}$ $E_h$ and $1\cdot10^{-3}$ $E_h\cdot\alpha^{-1}$, respectively, during the optimization process. Additionally, the optimization was carried out using the ALPB[4] water model, and GFN2[5] was chosen as the tight-binding model for all subsequent steps.

The optimized structure was used for a conformational search using CREST,[6,7] where iMTD-GC sampling in water was performed. The resulting conformer ensemble was narrowed down by performing energetic sorting using CENSO[8] (Parts 0-3 enabled). Again, the ALPB water model was for implicit solvation. The energy thresholds for the first three stages were set to 4.0 $kcal\cdot mol^{-1}$ (Part 0), 3.5 $kcal\cdot mol^{-1}$ (Part 1), and 2.5 $kcal\cdot mol^{-1}$ (Part 2), whereas a Boltzmann population cutoff of 99% was applied in the final stage (Part 3) instead. The respective functionals and basis set used were: b97-d3[9] and def2-SV(P)[10] (Part 0), r2scan-3c[11] (Parts 1 and 2), wb97x-d4[12] and def2-TZVPP[10] (Part 3), according to the defaults suggested by the CENSO developers. The number of conformers considered was limited to 200 to save computational resources.

Further information can be extracted from the CENSO configuration file supplied with the SI.

#### Electrostatic potential calculation

The electrostatic potential surface for each conformer of the sorted conformer ensembles was calculated by generating a Gaussian[13] input file using antechamber.[14] Redundant geometry optimization was avoided by modifying the input files to not include keywords associated with geometry optimization before execution of Gaussian. The HF/6-31G* level of theory was used for ESP calculation,[15,16] and the Merz-Singh-Kollman charge analysis scheme was implemented using 10 layers and 17 grid points per unit area.[17,18] The resulting output files were used for electrostatic parametrization as described in the main text.

#### Abbreviations

ALPB, Analytical linearized Poisson-Boltzmann; ANCopt, Approximate normal coordinates rational function optimizer; CENSO, Commandline Energetic Sorting; CREST, Conformer-Rotamer Ensemble Sampling Tool; ESP, Electrostatic Potential; GC, genetic structure crossing algorithm; GFN, geometries, frequencies and non-covalent interactions; HF, Hartree-Fock; MTD, meta-dynamics; PDB, Protein Data Bank; XTB, Extended Tight Binding;
